# A spatially resolved single cell proteomic atlas of Small Bowel Adenocarcinoma

**DOI:** 10.1101/2025.02.01.634535

**Authors:** Zeynep Dereli, Behnaz Bozorgui, Maga Sanchez, Nicholas Hornstein, Guillaume Thibault, Huamin Wang, Gordon B. Mills, John N. Weinstein, Michael J. Overman, Anil Korkut

## Abstract

Small bowel adenocarcinoma (SBA) is a rare malignancy marked with a poor prognosis. The cellular and proteomic heterogeneity within the tumor immune microenvironment (TIME) of SBA is a likely driver of prognosis, disease progression and response to therapy. We have addressed a major gap in knowledge of the TIME in SBA using highly multiplexed, protein imaging of the SBA tumor-immune ecosystem generating a comprehensive, single-cell level, spatial, proteomic atlas of TIME in > 600,000 cells from 136 tumor and matched normal samples from clinically and genomically annotated SBA patients (N=37). The SBA TIME Atlas informs on spatial distribution and interactions of tumor-intrinsic processes, diverse immune cell types, immune checkpoints, and vascularization. Enrichment of proliferating epithelial tissue, stem cells, and likely pro-tumor immune signatures in the tumor niche is contrasted by the representation of naïve and early-effector T-cells in the epithelial compartments of adjacent normal tissues. Epithelial-stem-immune cell spatial architectures within tumor and matched-normal niches were strongly associated with patient survival, suggesting malignancy is driven by spatial architecture beyond the tumor microenvironment. The blueprints of therapeutically actionable immune checkpoints at the interface between epithelial and microenvironmental T-cells as well as macrophages have established a guideline for precision immunotherapies tailored to the TIME composition in SBA. We expect that this SBA atlas will contribute to a deeper understanding of the immune contexture in this rare disease as well as other gastrointestinal cancers and help guide future precision immune-oncology strategies.

## Introduction

Despite marked advances in other gastrointestinal (GI) malignancies, deeper understanding of small bowel adenocarcinoma’s (SBA) biologic underpinnings remains nascent [1]. This lack of improvement can largely be traced to the rarity of small bowel adenocarcinoma and is reflected in continued poor outcomes for this patient population, Surprisingly, although the small intestine represents approximately 75% of the length and 90% of the surface area of the alimentary tract, the incidence of SBA is 50-times less than the more commonly occurring large bowel adenocarcinoma (colon cancer) [2–4]. A better understanding of the pathophysiology of SBA may also lead to critical insights into the more commonly occurring proximal counterpart, colon cancer. Though a multitude of hypotheses have been proposed, such as the rapid transit time in the small bowel, lower relative bacterial load, more alkaline environment, and greater lymphoid infiltrate in the small-intestinal mucosa, the explanation for the dramatic difference in incidence remains unknown [1].

Recent molecular analyses of SBA have demonstrated an unique genomic profile, the most pronounced difference from other GI adenocarcinomas being its rate of APC gene mutation (27%) as opposed to 76% for colon and 8% for gastric cancer [5]. SBA also has known differences from colon cancer in both stage distribution, metastatic predilection, and survival [6–8]. In a recent National Cancer Database stage-stratified comparison of survival between SBA and colon cancer, there was worse overall survival for SBA, even in Stage I patients with comparable lymph node sampling [8]. Although a small proportion of SBA are related to Lynch Syndrome or Celiac disease, the majority of SBAs are sporadic in etiology with no clear environmental or genomic risk factors [4].

Despite the known role of immune surveillance in the small intestinal mucosa, no comprehensive effort to understand the tumor-immune microenvironment (TIME) of SBA has been conducted. Limited studies have characterized the CD8 T-cell infiltration and PD- L1 expression within SBA [9–11]. One clinical trial investigated the efficacy of anti-PD1 pembrolizumab therapy in 40 SBA patients [12]. However, the trial demonstrated limited activity with a 3% response rate in patients with proficient mismatch repair SBA, further supporting the need for an improved understanding of the immune contexture of this rare cancer^4^.

The highly multiplexed imaging method, cyclic immunofluorescence (cycIF) is particularly useful for spatial mapping of the tumor immune contexture in single cell resolution [13]. Using the cycIF method, we generated a comprehensive spatial single cell proteomic atlas of SBA TIME to examine the relations between diverse clinical variables with immune and tumor parameters in both tumor and adjacent normal (originating from the same surgical sample with the matched tumor) tissue. Through analysis of the atlas, we found that the spatial distribution of immune activity, mesenchymal and endothelial cells in tumor and adjacent normal niche define the prognosis of small bowel adenocarcinoma. While the tumor niche is enriched by proliferating epithelial lineage cells and stem/mesenchymal characteristics, the normal niche is enriched with a population of active T-cells. Strikingly, spatial enrichment and interactions between early effector and naïve CD8+ T-cells with endothelial, mesenchymal/stem and epithelial cells in the normal niche, and macrophages with B-cells in the tumor niche are strong predictors of patient survival. Additionally, immune checkpoint receptor-ligand pairs are spatially enriched on Ki67+ epithelial and T-cells at the tumor normal interface providing opportunities for precision immunotherapy strategies in a significant fraction of patients. The presented spatial biology resource will aid in the development of therapeutic, clinical, and immunological characterization of this rare but devastating cancer type.

## Results

### A spatial atlas of tumor-immune interface in SBA in single cell resolution

We profiled the spatial distribution of markers associated with different cell types including epithelial, vascular, stem/mesenchymal, and immune cells, therapeutically actionable immune checkpoints, and various cancer hallmark proteins (cell signaling, proliferation, DNA repair, growth, vascularization). We used the cycIF method with validated antibodies that are conjugated to AlexaFlour antibodies on a clinically annotated tissue microarray (TMA) composed of 136 samples from 37 patients (**Figure 1A**, **Suppl. Figure 1**, see Methods for antibody information and validation approach). After quality control and filtering steps to eliminate samples with poor staining or tissue loss, the resulting resource included positional and expression data from > 600,000 cells on 111 samples of 35 patients mapped with 39 unique antibodies (**Figure 1B-D**). An analysis of tissue dropouts due to tissue losses demonstrated that we had readouts from > 400,000 cells with 36 protein markers and > 200,000 for the complete 39 antibody panel (**suppl. Figure 1**). We quantified the protein expression and location for each cell and marker using our computational analysis pipeline for data processing, segmentation, and protein quantification (see methods).

**Figure 1.**
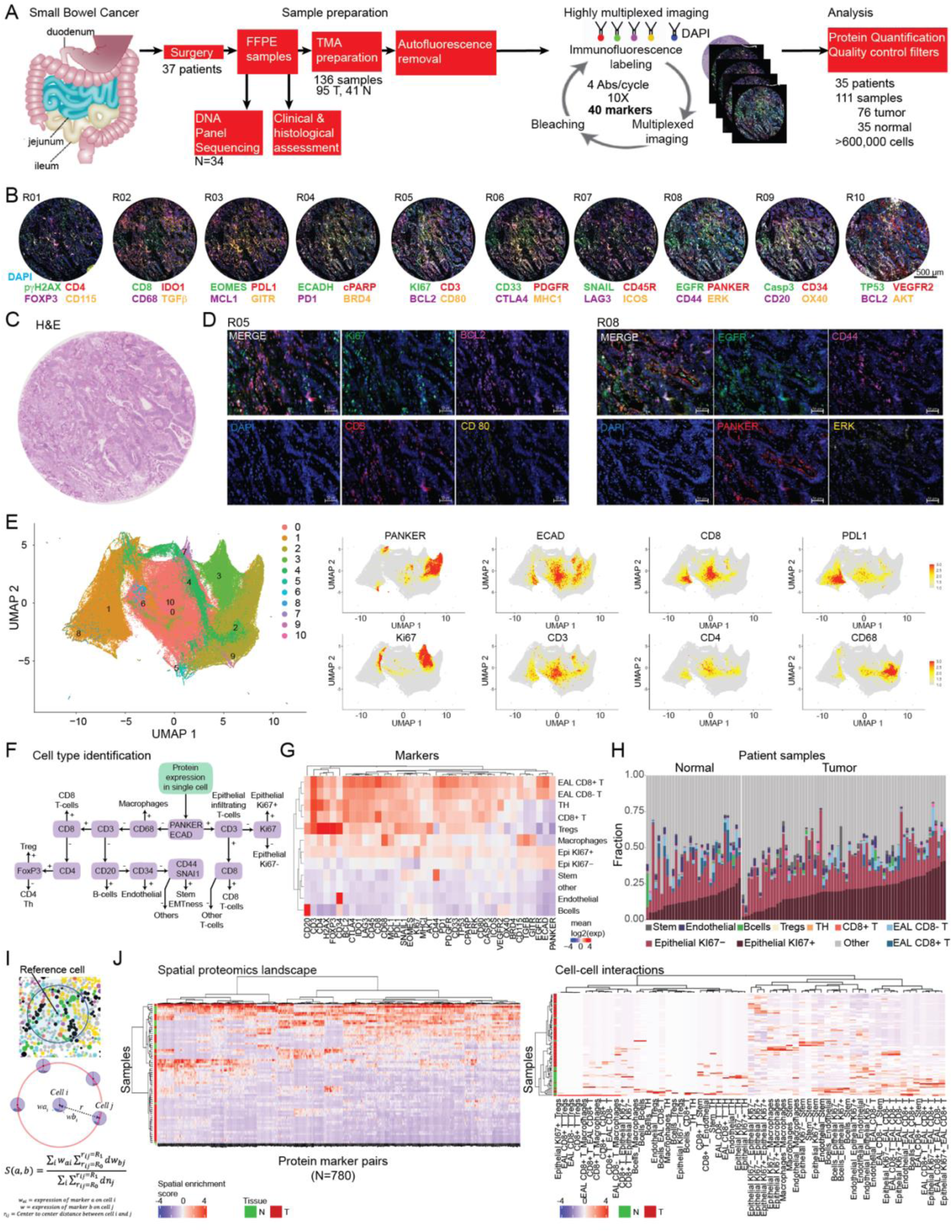
**The atlas of tumor-immune interactions in SBA**. **A.** Sample acquisition and preparation from surgery samples, cycIF-based spatial proteomics data collection and data analysis scheme. In cycIF, after an initial autofluorescence step, samples are stained with 4 conjugated antibodies per cycle with distinct excitation wavelengths (488, 555, 647, 750 nm). A fifth dye (e.g., DAPI) is used to stain nuclear DNA to locate cell nuclei so that signals from successive cycles can be compared within the same framework. After staining, samples are imaged using fluorescence illumination to interrogate target protein levels. After mild and rapid inactivation of fluorescent tags with H2O2 bleaching, samples are re-stained with another set of fluorescent antibodies. **B.** Up to 40 protein markers are measured on FFPE samples from SBA patients using the cycIF method. The raw data, the highly multiplexed tissue images are shown for one sample profiled for 40 markers in staining/bleaching/imaging 10 cycles (Suppl. figure 1 for the readouts from the complete tissue microarray). **C.** The H&E staining from the matched sample. **D.** A magnified view of the representative staining patterns on the ROIs from the same images in 1B in cycles 5 and 8. **E.** A cumulative UMAP (left) representation of single-cell level proteomics landscape quantified from the cycIF images. The expression values are log-transformed and Z-scaled. The UMAP covers both tumor and adjacent normal niches and involves > 180,000 cells. The distribution of select epithelial and immune markers as well as ICPs on the UMAP (right). **F.** The hierarchical scheme for cell type identification based on cell-type marker expression. **G.** The distribution of protein levels across the cell types. **H.** The fraction of cell types in each sample within the SBA cohort. **I.** The mathematical scheme for calculation of spatial relations between markers and cell types. The spatial enrichment score quantifies the expression enrichment of two protein markers in close proximity, and yet, likely in separate cells (1.66 μm < R < 10 μm). Similarly, for cell-cell interactions, the spatial enrichment score quantifies the proximal abundance of two cell types within the distance, R < 10 μm. The same equation is used for both marker and cell type spatial enrichment. For markers, the Z-scale log transformed expression levels quantified from cycIF images are used. For cell types, the annotations from the hierarchical cell type identification are used with binary quantification (1 if 1.66 μm <R <10 μm; 0 if R < 1.65 μm or R > 10 μm). **J.** The map of spatial interactions between proteomic markers (left) and cell types (right) on SBA samples as calculated using the approach in Figure 1I and analyzed with unsupervised hierarchical clustering method.

The resulting spatial proteomics atlas involves three information layers. Utilizing antibody combinatorics, the first information layer deconvolves the proteomic heterogeneity corresponding to distribution of protein expression levels across individual cells. The second information layer identifies cell types and major compartments including epithelial, immune, stem/EMT, and vascular cells. Finally, the third information layer is the spatial organization of protein markers and cell types. In the first layer, we enumerated the proteomic heterogeneity through annotation of proteins and expression levels in individual cells, combining tumor and adjacent normal tissues (**Figure 1E**). In the second layer, to analyze the composition and distribution of cell types, we implemented a hierarchical gating strategy based on well-established cell type markers (**Figure 1F-H**). This has led to the annotation of diverse epithelial, immune, endothelial, and stem/EMT cell types to > 300,000 cells while those cells that were silent for all cell markers remained either unidentified or associated with likely stromal cells based on a histological assessment. Third, we computed the pair-wise spatial enrichment score of protein markers and cell types to quantify cell-cell interactions in the tumor-immune microenvironment (**Figure 1I-J**). The spatial enrichment score is calculated for each sample as the expression-weighted density of marker or cell type pairs normalized with respect to the overall density of all cells in the sample. The spatial enrichment scores establish a rich resource for 835 interactions for proteins and cell types encompassing both tumor and adjacent normal tissue (**Figure 1J**). The resulting resource on tissue heterogeneity, cell type composition, spatial distribution and clinical characteristics of the patients will guide the analysis of tumor-immune interface atlas in SBA.

### Differential immune infiltration distinguishes the tumor and adjacent-normal niches

Using the spatial atlas, we analyzed the spatial heterogeneity in tumor and adjacent normal tissues. The spatial protein markers and cell type patterns led to clearly segregated clusters of normal and tumor niches suggesting architectural differences between the two (**Figure 1J**). Normal tissue is characterized by robust and recurrent spatial enrichment of heterologous cell-cell interactions while tumor tissue is sparse in recurrent spatial interactions suggesting the existence of a relatively disorganized tumor tissue structure.

First, we compared the differential enrichment of protein markers in the tumor vs. normal niches. Based on the statistical analysis of pseudobulk (sum of protein levels across individual cells within a sample) protein expression levels, we observed statistically significant enrichment of immune markers and checkpoints (CD8, CD68, CD3, PD1, PDL1, CTLA4, LAG3, IDO1, CD45R, CTLA4) as well as ERK signaling protein in normal tissue (Adjusted P value < 0.05 based on a two tailed t-test test with BH FDR correction) (**Figure 2A- B**). Within the tumor niche, the immunomodulatory proteins, TGFβ and GITR/TNFRSF18 as well as the proliferation marker, Ki67 are significantly enriched (Q-value < 0.05). The analysis of cell type distributions in tumor vs. adjacent normal niches suggested a highly significant enrichment of T-cells that spatially overlap with epithelial tissue (i.e., epithelial associated lymphocytes (EAL) including intraepithelial and lamina propria lymphocytes) within the normal niche (FDR Adjusted P value < 1e-4) **(Figure 2C,E-F)**. Endothelial, CD3+CD4+ Helper T (Th), and CD3+CD8+ T cells as well as macrophages were significantly enriched within the normal niche although not as strikingly as EAL CD8+ T cells (Adjusted P value <0.05). The tumor-harboring tissue, on the other hand, is significantly enriched with Ki67+ epithelial cell (Adjusted P-value < 1e-3) as well as CD44 or SNAIL expressing mesenchymal/stem cell populations (Adjusted P-value < 1e-2). Interestingly, the presence of EAL CD8+ T-cell in normal tissue is associated with a trend towards better survival (log rank P-value=0.12) (**Figure 2D**). In conclusion, the analysis suggests a significantly increased immune infiltration within normal tissue and a reciprocal immune depletion yet proliferating epithelial cells and stemness within the tumor niche.

**Figure 2.**
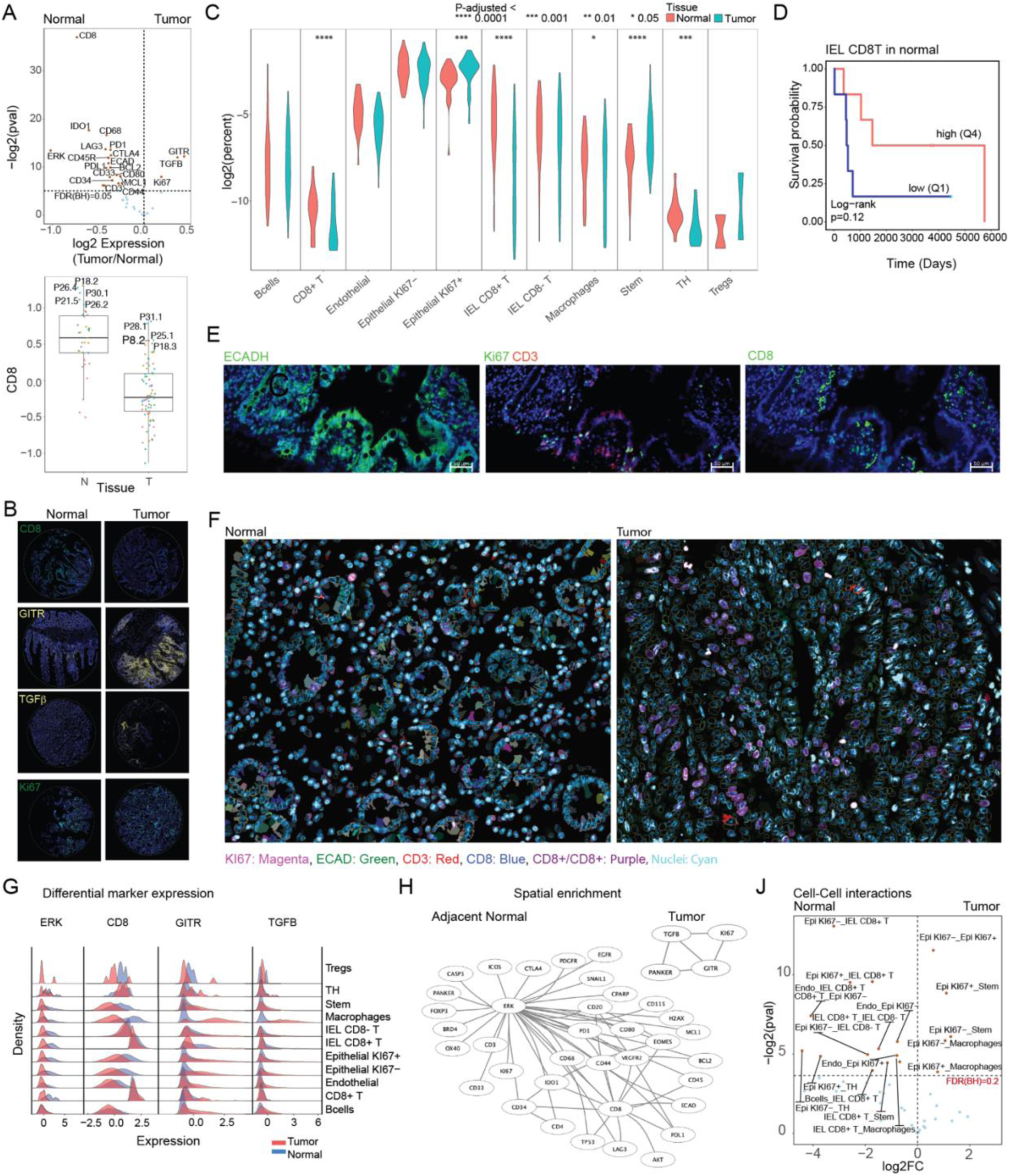
The immune microenvironment landscape in tumor-containing and tumor-adjacent (normal) tissue in SBA. **A.** Top: The differential enrichment of protein markers in tumor vs. tumor-adjacent normal niches. Bottom: The comparison of CD8 expression in tumor vs. normal microenvironment (right). The Q-values are based on a Wilcoxon test followed by FDR-correction with BH method. **B.** Representative images of differential expression patterns in matched tumor vs. adjacent normal tissues in select patient samples. **C.** The abundance of cell-types annotated through marker expression in tumor vs. tumor-adjacent tumor niches. The p-values are based on an unpaired two-tailed Wilcoxon-test (FDR-adjusted with Benjamini-Hochberg method) between samples corresponding to tumor-adjacent normal vs. tumor niches. **D.** The overall survival difference in patients carrying low vs. high abundance of CD3+/CD8+ epithelial infiltrated lymphocytes (EAL CD8+ T cells) in adjacent normal tissue. **E.** Representative raw images of E-cadherin, Ki67, CD3 and CD8 expression in epithelial tissue within the tumor-adjacent normal niche. **F.** Representative spatial feature maps showing the expression of E-Cadherin, Ki67, CD3, and CD8, calculated through segmentation and quantification of raw images from matched normal (Sample 81) and tumor (Sample 85) tissues from the same patient. The spatial features are generated by mapping the registered, processed, and quantified protein expressions on coordinates of individual cells. **G.** The single cell-level distribution of differentially expressed markers ERK, CD8, GITR, and TGFB across cell types in tumor vs. tumor-adjacent normal tissues. **H.** The network representation of protein markers that are spatially co-enriched in tumor-adjacent normal (top) and tumor (bottom) niches. The nodes are the proteomic markers enriched in the respective niche. The edges are constructed for spatial co-enrichments for absolute log2(FC)> 1.5 where FC: Fold change of mean in tumor vs. normal niches, P-value < 0.05 based on a two-sided t-test. **J.** The statistically significant spatial enrichments in tumor-adjacent normal vs. tumor tissues. FC: Fold change of mean in tumor vs. normal. The p-values are based on an unpaired two-tailed t- test.

Next, we mapped the protein markers to individual cell types with particular focus on differentially expressed proteins (**Figure 2G**, **Suppl. Figure 2**). Expression differences in ERK in the tumor vs normal tissue was reflected in the expression differences in T cells, suggesting potential ERK pathway activity downstream of T cell receptors and resulting T cell activation in adjacent normal tissue. Macrophages within the tumor niche express significantly higher TGFβ and GITR levels compared to macrophages in the normal niche, suggesting macrophage heterogeneity across niches. The spatial co-enrichment analysis of protein markers identified ERK and CD8 as hubs within the normal tissue such that a series of proteomic markers were enriched around cells consistent with the high expression levels of the two markers **(Figure 2H**). On the other hand, within the tumor niche, a network of proteins involving TGFβ, GITR, PanKeratin and Ki67 were spatially co-enriched across all samples. Finally, the spatial analysis of cell-cell interactions demonstrates that the tumor niche is highly enriched in interactions of epithelial cells with macrophages and stem/mesenchymal cells whereas the normal niche is dominated by interactions of T-cells with the epithelial cells (**Figure 2J)**. In conclusion, we observed mutually exclusive representation of cellular heterogeneity as well as spatial interactions between the tumor and adjacent normal niches.

### The Immune checkpoints spatially overlap with immune and epithelial cell interfaces

We analyzed the spatial distribution of therapeutically actionable immune checkpoints across diverse cell types and tissue niches; these targets included PD1, PD-L1, CTLA4, ICOS, LAG3, CD115/CSF1R, CD80, IDO1, and GITR/TNFRSF18. As demonstrated through an analysis of expression levels across cell types, the immune checkpoint expression was predominantly enriched in immune cells, particularly the T-cell populations in both tumor and adjacent normal tissue (**Figure 3A**). The T cells were characterized by high expression of PD1, PDL1, LAG3, CTLA4, and IDO1 while other ICPs were present at relatively low levels (**Figure 3B**). In macrophages, there was substantial variation between the tumor vs. adjacent normal niches with CD115 and GITR enrichment in the former and PDL1 and IDO1 enrichment in the latter. Treg cells were enriched with PDL1 interactions in both niches as well as moderate CTLA4 expression within the tumor niche.

**Figure 3.**
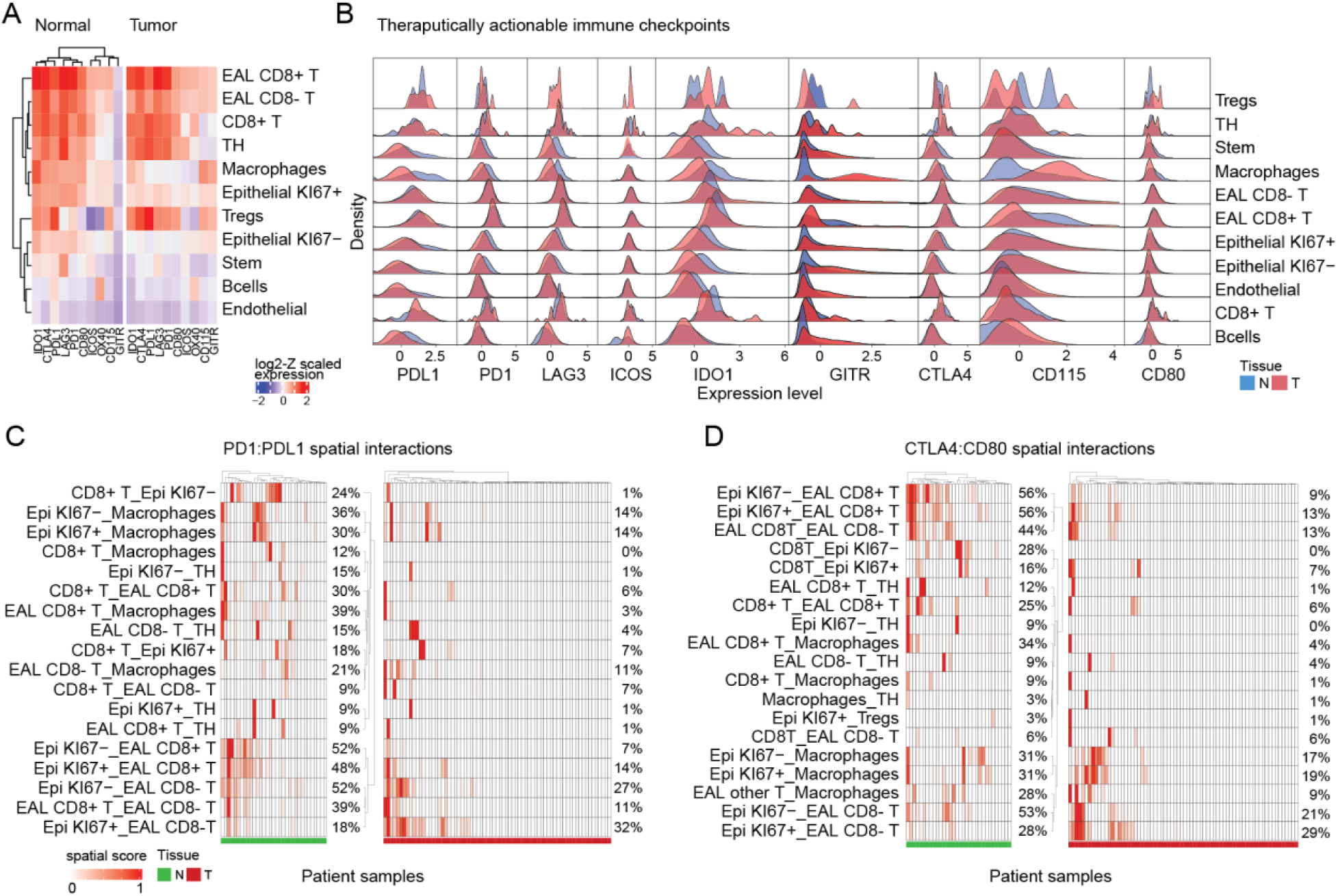
Spatial heterogeneity of immune checkpoints (ICPs) in the SBA ecosystem. **A**. A heatmap of ICP expression in diverse cell types averaged over tumor (left) and adjacent normal (right) samples. **B.** The histograms that demonstrate the heterogeneity and distribution of therapeutically actionable ICPs in individual cells from tumor (T, red histogram) and adjacent normal (N, blue histogram) samples. **C.** The immunoprint analysis visualizes the distribution of PD1:PDL1 (left) and **D.** CTLA4:CD80 interactions as represented by spatial co-enrichment (spatial enrichment score > 0) in specific cell type pairs across tumor and adjacent normal samples. The frequency of each spatial co-enrichment is given as percentiles on the right-hand site of the immunoprints. The green and red bars underneath the immunoprints indicate the <u>normal and tumor samples, respectively</u>

To model immunoregulatory interactions, we mapped the spatially proximal ICP receptor-ligand pairs between heterologous cells based on the spatial enrichment scores. We focused on the therapeutically relevant PD1-PDL1 (**Figure 3C)** and CTLA4-CD80 (**Figure 3D)** interactions which may inform precision immunotherapy decisions in future translational studies. The analyses led to quantification of the sample specific occurrence and population frequency of PD1-PDL1 and CD80-CTLA4 interactions (enrichment score > 0) in specific cell type pairs represented on “immunoprints” in similar spirit to widely used oncoprints [14]. Consistent with immune infiltration profiles (**Figure2**), both PD1-PDL1 and CD80-CTLA4 interactions were more frequent within the adjacent normal tissue. The potential PD1-PDL1 interactions are most frequently carried between the populations of epithelial cells, EAL T-cells and macrophages. Within the adjacent normal tissue, positive PD1-PDL1 enrichment scores were observed in 52% of samples on proximal CD8+ T and Ki67- epithelial cells, 38% of samples for proximal macrophages and Ki67- epithelial cells, and 36% of samples for proximal macrophages and CD8+ T cells. The most frequent PD1- PDL1 interaction between heterologous cell types was between the epithelial and CD8- T cells (32% for Ki67+ and 27% for Ki67- epithelial cells in proximity of CD8- T cells). Similarly, the CTLA4-CD80 co-enrichment was most frequent between proximal epithelial and CD8+ T cells in the adjacent normal (56% of normal samples) and epithelial and CD8- T cells within the tumor tissue (29% of tumor samples). The CTLA4-CD80 co-enrichment on Ki67+ epithelial and macrophage cells was substantially more frequent in adjacent normal compared to the tumor niche (31% vs. 19%). In conclusion, we observed a spatial co-enrichment of PD1-PDL1 and CTLA4-CD80 ICP receptor-ligand pairs, which indicates immunoregulatory interactions mediated between epithelial and T-cell as well as macrophage populations. The findings provide a quantitative account of ICP interactions across tumors of SBA patients and are consistent with increased immune infiltration within the tumor adjacent normal niche. The immunoprint analysis provides a framework for selection of precision immunotherapies tailored to recurrent ICP receptor-ligand interactions in specific cell types that are frequently in proximity.

### Heterogeneity of the epithelial-immune interface in tumor and adjacent normal niches

We addressed how immune heterogeneity in SBA ecosystem modulates disease progression through a comprehensive analysis of the T-cell and macrophage populations. Guided by a set of discriminant protein markers (CD3, CD8, CD4, EOMES, PD1, FOXP3), we identified the distribution of T-cell subtypes within the intraepithelial neighborhood where T-cell abundance is highest. Using marker information with single-cell resolution, we deployed a detailed gating algorithm for lineage separation (**Figure 4A**). The analysis led to the quantification of CD3+/CD8+ T-cells that fall into subgroups of early effector (PD1+ / EOMES-), late effectors (PD1- /EOMES+), exhausted/activated (PD1+ / EOMES+) and native T-cell groups (PD1- / EOMES-). We have also mapped the EAL Th (CD3+ / CD4+), Tregs (CD3+ / CD4+ / FOXP3+) and other CD3+ T-cells for which we did not detect CD4 or CD8 expression (double negative). While the overall enrichment of T-cells within the adjacent normal tissue is consistent with our analysis demonstrated in Figure 2, the refined analysis including T-cell subpopulations makes clear that this enrichment is driven by the significantly increased early-effector and naïve CD8+ T-cells (**Figure 4A**). The enrichment of early effector and naïve CD8+ T-cells suggests an active immune state in the adjacent normal tissue at least in comparison to the tumor niche. An immunoprint analysis of T cell spatial interactions in the adjacent normal niche reveals a cluster of patient samples in which early effector (CD3+/CD8+/PD1+), naïve (CD3+/CD8+), EAL Th (CD3+/CD4+), double negative T cell populations, macrophages and Ki67- epithelial cells are recurrently (> 40% of samples) in proximity with each other (**Figure 4B**) (spatial enrichment score > 0, **Figure 1I**). A mutually exclusive second cluster is characterized by interactions of exhausted T cells (CD3+/CD8+/PD1+/EOMES+), late effectors (CD3+/CD8+/EOMES+), macrophages and Ki67+ epithelial cells. In the tumor niche, the Th, double negative T-cell, Ki67+ epithelial cell interactions were observed in ∼20% of the samples. Within the tumor niche, weak interactions between Ki67+ epithelial and double negative T-cells were observed in 38% of the samples.

**Figure 4.**
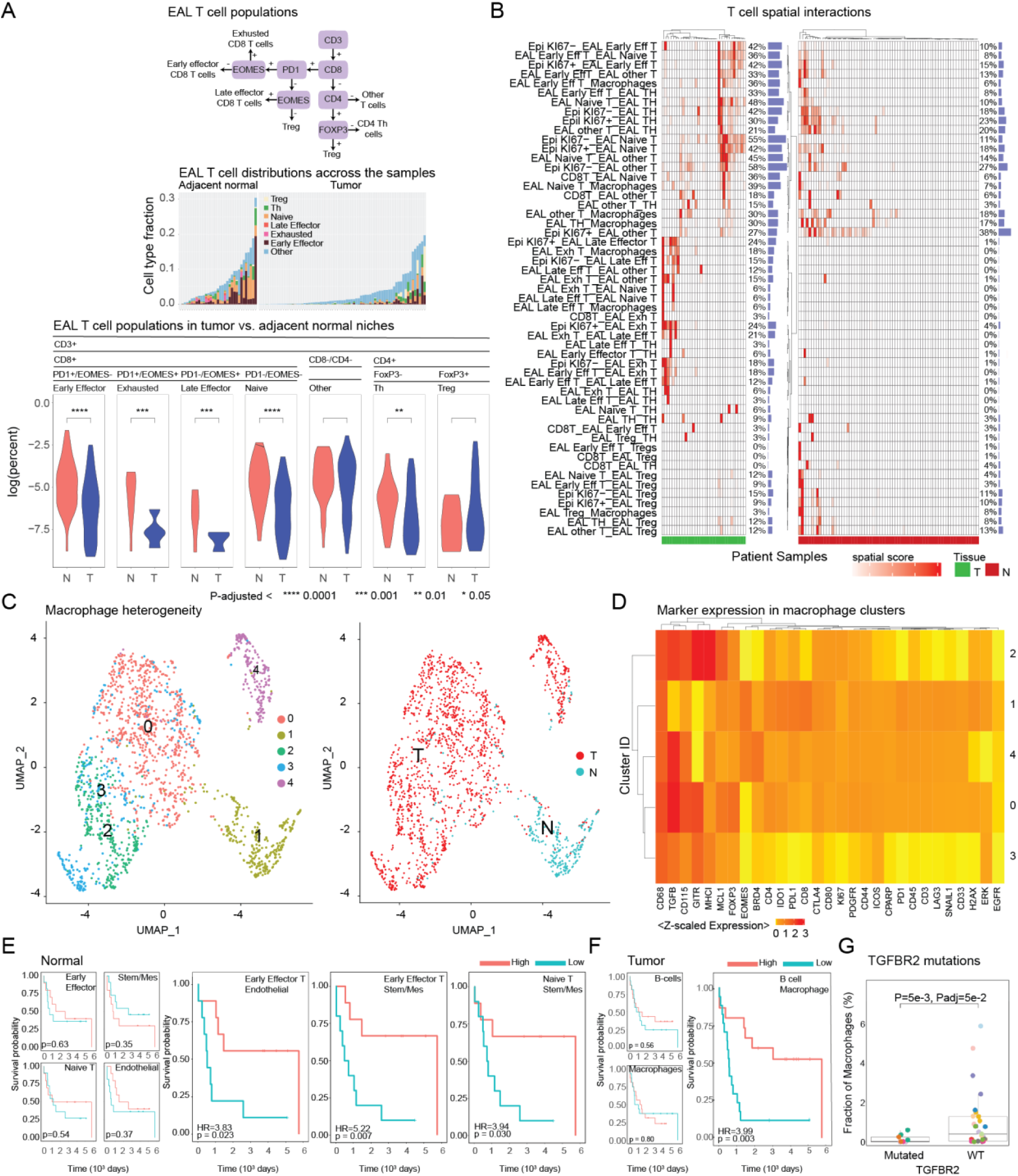
Spatial heterogeneity of T-cell and macrophages in the SBA ecosystem. A. The distribution of T-cell subpopulations in SBA. The hierarchical scheme for intraepithelial T-cell subpopulations (top left). The distribution of T-cell subpopulations across tumor and adjacent normal samples (top right). The comparison of T-cell populations in tumor vs. normal samples (bottom). The P-values are based on a 2-tailed t-test followed by FDR-adjustment with Benjamini- Hochberg method. **B.** The interactions of T-cells with all cell types as quantified by cell-cell spatial enrichment scores across the SBA tumor and adjacent normal tissue samples. **C**. Left: UMAP representation of macrophage heterogeneity. The macrophages (N=1840) with complete protein marker expression data are included in the analysis. covering the macrophages based on protein marker expressions. Right: The macrophages in tumor vs. adjacent normal tissue fall into distinct clusters on UMAPs. **D.** Protein marker expression profiles of each macrophage cluster averaged over all cells in the respective cluster. **E.** The impact of T-cell interactions within adjacent normal tissue and **F.** T-cell populations in tumor domain on patient survival as demonstrated by Kaplan- Meier (KM) curves. The cohorts are stratified as “high” vs. “low” group based on the median of the spatial enrichment scores. The p-values are based on a log rank test. On the left side, the impact of individual cell type abundance (small-sized KM-plots) on patient survival are demonstrated with KM-curves with a patient stratification based on median abundance of each cell type. **G.** The association of TGFBR2 mutations with macrophage abundance in tumor microenvironment. The p-value is based on an unpaired two-tailed t-test between the TGFB2R2 mutated vs. wild type cells. The adjusted P-values are based on FDR-corrected using the BH- <u>method.</u>

Next, we studied the heterogeneity of CD68+ macrophage cells. K-means clustering [15] of the single cell protein expression data from 1840 macrophages across all samples (N=111) yielded 5 distinct clusters as represented in the UMAP analysis (**Figure 4C**). Strikingly, the macrophages from the adjacent normal tissue were populated in a single and isolated cluster (cluster 1) on the macrophage UMAP whereas the macrophages from the tumor niche are spread across four clusters (clusters 0,2-4). The marker expression analysis demonstrated that all clusters carry high CD68 expression as expected while TGFβ, CD115 and GITR are represented at high levels in the tumor niche macrophages (**Figure 4D**). The differential analysis of each cluster suggests that the tumor niche clusters were differentially enriched with MHC1, MCL1, CD115 (cluster 3); TGFB, FOXP3, CD115 (cluster 0); BRD4 (cluster 4) (**Suppl. Figure 4**). Cluster 0 was silent for all markers in the differential expression analysis. The expression of TGFβ, MHC1 and CD115 within the tumor niche is consistent with enrichment of tumor promoting M2-like macrophages [16–18]. The normal niche-associated cluster 1 is particularly depleted in TGFB, GITR, MHC1, and CD115 while relatively enriched for a large series of markers including CD80, suggesting a possible representation of anti-tumor M1-like macrophages. While PD1, EOMES, and CD8 are not at high levels in the macrophages of the normal niche, they are at high levels compared to macrophages in the normal niche (**Suppl. Figure 4**). Overall, our comprehensive analyses of T-cell and macrophage heterogeneity across the SBA ecosystem suggest the presence of a more active immune state within the adjacent normal tissue characterized by increased naïve as well as early effector cells, while the tumor niche is marked with tumor-promoting macrophage populations.

### Spatial distributions of immune cells modulate patient survival

Motivated by the differential enrichment of immune cells across tumor and normal niches, we analyzed relationships of cell-cell interactions with overall survival. We stratified patient groups into low and high groups for each parameter (i.e., cell type abundance and spatial enrichment scores for interactions) based on median values. The survival differences between the low vs. high group were analyzed with Kaplan-Meier methods and log-rank statistics. There were striking differences in patient survival based on spatial enrichment scores within both tumor and normal niches (**Figure 4E-F** and **Suppl. Figure 4**). In the tumor niche, B-cell and macrophage proximal co-enrichment was the strongest prognostic predictor where high co-enrichment correlated with better survival (P-value=3e-3, Hazard-Ratio (HR)=3.99). In the normal niche, EAL naïve CD8+ T, early effector CD8+ T, endothelial and stem/mesenchymal cell interactions were most strongly correlated with patient survival (P-value < 0.03, 3.8 > HR for low vs. high pairwise interactions). Within the normal niche, we also observed several strong trends involving interactions of EAL T cell groups with epithelial, B-cell and macrophages such that high spatial co-enrichment predicted better patient survival for each case (log-rank P value< 0.2, **suppl. Figure 4**). While T cell subtype abundances were low in the tumor niche, we observed that spatial enrichment between the EAL Naïve and early effector T cell populations against epithelial, endothelial and stem cells were associated with better survival in contrast with the adjacent normal tissue. This observation possibly suggests a balance between the tumor and adjacent normal tissue such that patients have mutually exclusive EAL T-cell enrichment in tumor vs. adjacent normal tissues leading to the opposite survival correlations. An exception to this pattern was the EAL Th and endothelial interactions within the tumor niche, which correlated with better survival. In contrast to spatial enrichment patterns, the abundance of immune cells had moderate to no impact on patient survival (All P-values > 0.05 based on a log-rank test). In conclusion, the spatial interactions are relevant and suggest a tumor ecosystem state that modulates patient survival through interactions carried by pro-tumor macrophages and active (i.e., naive, or early effector) anti-tumor CD8+ T cell populations in tumor and adjacent normal niches, respectively.

### TGFBR2 mutations are associated with macrophage depletion in TIME

We investigated the associations between frequent mutations in SBA tumors with the immune parameters. We utilized a previously published targeted-exome sequencing data for which matched cycIF data is available for 18 patients [19]. Due to the limited overlap between the cycIF and mutation data, we focused on the most frequently mutated genes to gain statistical power. Our analyses involved *TP53* (8 out of 18 patients with mutated tumors), KRAS (5/18), PTEN (3/18), PIK3CA (3/18), ARID1A (6/18) and TGFBR2 (5/18) (**Suppl. Figure 4**). We merged the PTEN and PIK3CA mutated groups to establish a PI3K/AKT pathway mutated group. Next, for each gene, we compared the abundance of various cell types in mutated vs. unmutated samples. Across all the cell types and genes, the only statistically significant association is observed as the depletion of macrophages in TGFBR2-mutated tumor samples (t-test adjusted P value =0.05, log2 fold change =-2.70 for mutated vs. wild- type) (**Figure 4G**). No other cell type was detected as significantly enriched for any cell type in tumor or adjacent normal niches. Considering the observed enrichment and previously defined roles of TGFβ signaling in tumor associated macrophages, further studies are warranted for functional consequences of TGFBR2 mutations in the context of macrophage- tumor interactions in SBA.

### A clinically annotated spatial proteomics resource of immune-tumor interactions of SBA

We established a resource that links the spatial TIME and clinical variables in SBA tumors (**Figure 5**). The resource involves seven clinical variables including disease stage, anatomic location, ancestry, MMR-status, tumor differentiation, patient age and adenocarcinoma status (**Figure 5A**). The clinical variables are linked to abundance and spatial distribution of 39 unique protein markers and 11 cell types inferred from marker expressions. The resulting proteomics-based resource covers the differential enrichment of expression and spatial parameters for markers and cell types in > 300,000 cells within tumor and normal niches from clinically annotated samples.

**Figure 5.**
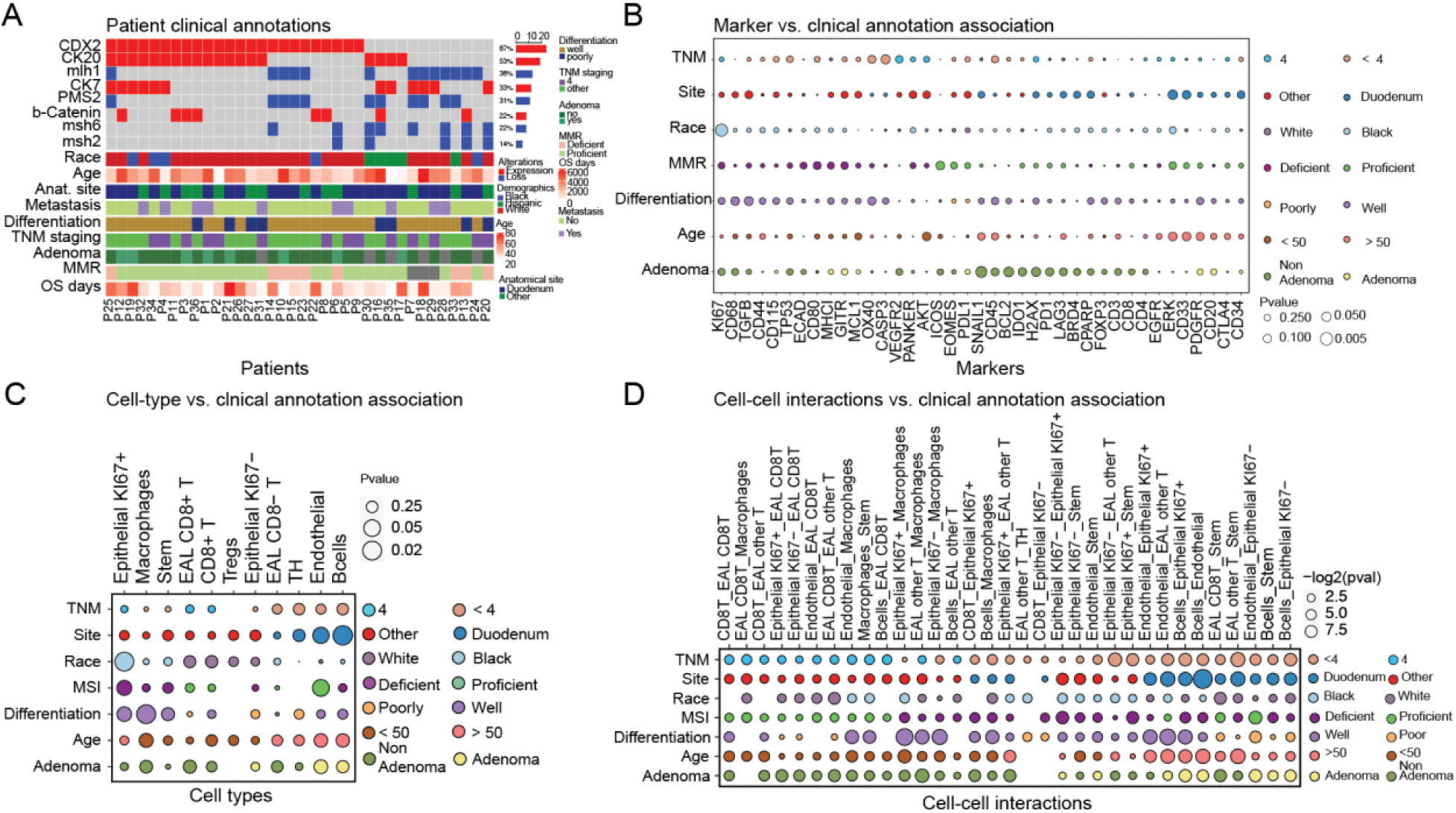
The association of spatial tumor-immune markers with clinical annotations. **A.** The clinical annotations of the patients in the SBA cohort. **B.** The differential expression of protein markers with respect to clinical annotations in tumor niche. The color-codes represent the higher expression in a specific clinical annotation (e.g., blue: duodenum, red: other sites). The circle sizes correspond to the statistical significance of the expression differences between clinically distinct groups based on an unpaired, two-sided t-test. **C.** The differential abundance of cell types with respect to the clinical annotations in tumor tissue. The color coding and statistical analysis are as in 3B. The cell type annotations are based on the hierarchical analysis in figure 1H. **D.** The differential analysis of spatial enrichment scores for cell-cell interactions with respect to the clinical annotations. The color coding and statistical analysis are as in figure 5B.

Overall, we mapped 5758 enrichments for abundances (N=266 for markers, N=77 for cell types) (**Figure 5B-C**) and spatial relations (N=5187 for markers, N=238 for cell type pairs) against clinical variables (**Figure 5D, Suppl. Figure 5**). The most striking associations that we detected within the resource are as follows. In protein expression and cell type profiles, we observed a significant enrichment of Ki67 expression (P-value < 5.0e-3) **(Figure 5B)** and Ki67+ epithelial cells (P-value < 2.0e-2) **(Figure 5C)** in the samples from Black patients compared to the non-Black group. This finding is consistent with the worse prognosis and higher Ki67 expression observed in Black patients across multiple cancer types [23]. We observed a significant endothelial cell enrichment in tumors from the Duodenum site compared to other sites (P-value < 5.0e2) **(Figure 5C).** The duodenum was also enriched with spatially proximal endothelial and EAL T-cell populations suggesting increased inflammation at this site(P-value < 5.0e-2) **(Figure 5D)**. The tumors with Mismatch repair proficient (pMMR) tumors were relatively enriched with endothelial cells while samples with MMR deficient (dMMR) tumors had enrichment of epithelial cells (P-value < 5.0-e2) **(Figure 5C)**. Well differentiated tumors carried increased macrophage populations as well as increased spatial proximity of the epithelial tissue and macrophage cells (P-value < 2.0e-2) **(Figure 5B-C)**. Consistent with the macrophage enrichment, we detected spatial interactions between macrophages and a variety of cell types including Ki67+ epithelial cells, B-cells, and T-cells in the differentiated tumors (P-value <5e-2) **(Figure 5D)**. While we cite the strongest associations, the spatial-clinical resource is highly rich and can be mined considering specific hypotheses, and potentially can provide a framework for building similar spatial resources for other cancer types in the future.

## Discussion

We present a comprehensive analysis of proteomic and cellular spatial heterogeneity within the SBA tumor ecosystem. The resource is based on cycIF spatial proteomic profiling of > 600,000 individual cells in the tumor and normal niches of clinically annotated surgical samples from 37 patients. The profiles cover diverse protein markers of epithelial, immune, mesenchymal, and vascular contexture, oncogenic processes (e.g., cell proliferation, apoptosis, DNA repair) as well as therapeutically actionable ICPs. Deep computational analyses of the spatial data revealed for the first time the multicellular contexture in the tumor and normal niches. We demonstrated that the spatial analysis of the SBA ecosystem can map tissue architecture and heterogeneity, immune activation states, therapeutically actionable ICPs, and spatial markers of prognosis in both tumor and associated normal tissue.

A comparison of spatial patterns in tumor and adjacent normal tissue showed normal tissue is globally more structured with recurrent spatial features across samples (**Figure 1H**). The contrast between tissue architecture in tumor vs. normal tissue may allow for automated diagnostic approaches particularly when coupled with emerging artificial intelligence methods for histology and warrants further studies in other cancer types. Motivated by this global architectural difference in tumor vs. normal niches, we focused our analysis on the differences in their cellular and molecular characteristics. We observed a significant enrichment of anti-tumor microenvironment within the adjacent-normal niche compared to the tumor niche. The normal niche was significantly enriched with various immune markers and cell types particularly the EAL CD8+ T-cells. A deeper analysis of the T-cell subpopulations suggested the EAL T-cells in the normal niche also corresponded to active naïve and early-effector states within the spatial neighborhood of the epithelial tissue, consistent with the antitumor immune composition. On the other hand, the tumor tissue was marked with spatial enrichment of stem/mesenchymal and macrophage cells within the Ki67+ epithelial structures. While the overall abundance of CD68+ macrophages did not differ significantly between the tumor and adjacent normal tissue, the macrophage types were heterogeneous as demonstrated on UMAPS and differential analyses (**Figure 4C-D, suppl. Figure 4**). The macrophages located in the normal niche were enriched within a single cell cluster on the UMAP consistent with a CD80+ M1-like macrophage dominance supporting a likely anti-tumor immune environment [24]. The macrophages in the tumor-niche, however, were distributed to three clusters, with high expression of CD115, TGFβ, and GITR that were supporting a likely pro-tumor M2-like polarization with spatial enrichment around the epithelial domain. Contrasting with the normal tissue macrophages, the tumor niche macrophages were significantly depleted in the presence of TGFBR2 mutations. The macrophage depletion within the TGFBR2 mutated tumor environment is consistent with the well characterized roles of TGFβ signaling on M2-like polarization [20]. The tumor-niche was also enriched with Ki67+ epithelial and stem/mesenchymal cell populations, consistent with the malignant phenotype. Overall, our analyses suggested an anti-tumor barrier within the adjacent normal tissue with high abundance and likely active CD8+ T cells, while the tumor core already reflected immune, mesenchymal/stem and proliferative characteristics consistent with malignancy.

The striking association between patient survival and multicellular neighborhood patterns as compared to overall cell numbers, demonstrate the clinical relevance of spatial contexture. Specifically, spatial enrichment patterns in the normal niche involving EAL early effector or naïve CD8+ T cells with endothelial, mesenchymal, and epithelial cells as well as those in the tumor niche involving B-cells with macrophages predicted significantly better patient survival. The association of patient survival with spatial patterns points to two important conclusions. First, the adjacent normal tissue is an active component of the tumor ecosystem as the spatial composition of active immune cells in the adjacent tissue associates with disease progression. Second, disease progression is mediated by the spatial enrichment patterns and likely interactions but not mere abundance levels of immune markers. This observation provides a rationale for spatial profiling to understand disease progression and potentially guide clinical decision making with spatial data. In conclusion, the pattern of co-existing spatial interactions in both tumor and normal tissue suggests presence of a spatial immune signature and activity that can predict patient outcomes. Of particular importance, the spatial composition of the adjacent normal tissue is a significant contributor to this prognostic signature. The active immune composition in the normal compartment is strongly associated with good patient survival, suggesting an “immune shield” that prevents spread and progression of the disease.

We devised the immunoprints to provide a proof-of-concept precision immune- oncology guide. The immunoprints quantify the immune checkpoint receptor-ligand pairs that are co-enriched in neighborhoods of immune and other cell types. Using the immunoptrints, we developed an analysis of PD1:PDL1 and CTLA4:CD80 co-enrichment patterns. Both receptor-ligand pairs were more recurrent within the adjacent normal tissue consistent with the immune cell and marker enrichment patterns. Both PD1: PDL1 and CTLA4:C80 spatial enrichment was most recurrent in the epithelial-EAL T cell neighborhoods independent of the Ki67 and CD8 status followed by the epithelial-macrophage neighborhoods. We observed three clusters for PD1:PDL1 enrichment neighborhoods. The first cluster involves epithelial-EAL T cells. The second cluster is marked by epithelial-macrophage or epithelial-CD8+ T (not infiltrated to epithelial tissue) interactions and third cluster is silent for any enrichment. The CTLA4:CD80 interactions, however, were represented in two clusters where the first cluster was enriched with strong epithelial- EAL T-cell interfaces and the remainder were either silent or marked with weak enrichment of macrophages or CD8+ T cells with the epithelial tissues. Overall, the immunoprints provide an analysis tool for defining patient groups based on spatial enrichment neighborhoods and may guide immunotherapies tailored to recurrent neighborhood patterns in tumor ecosystems.

Despite providing a deep insight into the immune and microenvironment structure of SBA, our study carries important limitations. First, the selection of 40 markers, although pointing to a high degree of multiplexing, is limited in covering all relevant interactions particularly for cell typing. We expect a majority of the “dark matter” in the cell type analysis corresponds to stromal tissue such as fibroblasts. While this is an important limitation as fibroblasts can mediate various immune and malignant phenotypes, it can be resolved with inclusion of stromal markers such as vimentin or histopathological annotations with H&E staining. Second major limitation is in precision of potential segmentation. While we have a robust segmentation algorithm, the complexities in the 3-dimensional cell morphologies particularly within dense regions such as epithelial tissue prohibits definition of cell boundaries with absolute certainty. For example, it is nearly impossible to map the membrane markers to the individual cells within tightly packed multicellular regions. Therefore, we have quantified the enrichment of molecular interactions (e.g., ICP receptor-ligand pairs) within the cellular boundaries without assigning each receptor to an individual cell. This limitation can be resolved with improved segmentation algorithms that use more advanced machine learning approaches that are trained with rich expression and multicellular interaction patterns. Third limitation is the dropouts in cellular readouts due to tissue damage during the tissue staining/destaining cycles with mild H2O2 treatment. While we were able to obtain molecular data from nearly 400,000 cells out of 600,000 until cycle 9, the coverage dropped to 200,000 cells in the last cycle. We expect the tissue damage can be overcome with new antibody technologies involving nucleotide conjugated dyes that can be stripped without major damage to the tissue. Fourth, the patients were treated with a variety of different therapies. Future studies of cohorts from clinical trials will be necessary to link the observations herein with specific therapy approaches.

The spatial atlas of SBA ecosystem is a highly rich resource to generate hypothesis to further investigate this rare disease and develop treatment strategies to improve patient survival. We expect future comparative studies with other gastrointestinal cancers as well as normal tissue samples will also benefit from the resource to identify how the tissue ecosystems and immune interactions vary across the gastrointestinal tract. Our study demonstrates how normal tissue can mediate immune and other features that directly impact disease progression and patient survival. It is likely that the future clinical/translational studies will need to account for the immune impact in the normal tissue to identify patient cohorts that will benefit from immunotherapy with deeper and more durable responses. Overall, we expect the spatial atlas of SBA with newly identified tumor-immune-mesenchymal cell interactions will contribute to improved treatment strategies for this rare disease while supporting similar research for more common GI cancers such as colorectal, liver, and pancreatic cancers.

## Methods

### Patient samples and construction of SBA TMA

The samples originate from patients undergoing treatment at the University of Texas MD Anderson Cancer Center (MDACC). The clinical annotations include demographics, cancer treatment history, stage, tumor differentiation, mismatch repair status, and survival for each patient (**Suppl. Table 1**). Tissue microarrays (TMAs) were constructed to include all clinically and histologically annotated samples, tumors (N=95) and adjacent normal tissue (N=41) (**Suppl. Figure 1**). [21] The hematoxylin and eosin (H&E) stained sections of the tumor and matched normal small bowel from 37 patients who underwent resection at our institution were reviewed. Representative formalin-fixed paraffin embedded tissue blocks of the tumor and matched normal from each patient were selected. Three 1.0 mm cores from each tumor and two 1.0 mm core of the matched normal small bowel tissue were used for the TMA construction. The TMA was constructed as previously described using a tissue microarrayer (Beecher Instruments, Sun Prairie, WI) [22]

### Reagents and antibodies

All conjugated primary antibodies and secondary antibodies with dilution, vendor and conjugation information are listed in **Suppl. Table 2**. 10mg/ml Hoechst 33342 stock solution was purchased from Life Technologies (Cat. H3570). 20X PBS was purchased from Santa Cruz Biotechnology (Cat. SC-362299). 30% hydrogen peroxide solution was purchased from Sigma-Aldrich (Cat. 216763). PBS-based Odyssey blocking buffer was purchased from LI-COR (Cat. 927-40150). CyCIF Heat induced epitope removal (HIER) solution was prepared as follow: Tri-Sodium citrate: 2.94g Distilled water: 1 L. Adjust to pH 6 adding 1N HCL (0.821ml of HCL to 10mL DH2O) and 0.5mL of Tween 20 was added and mixed. Buffer was stable for 3 months at room temperature or longer if stored at 4°C. Tris-EDTA Buffer (10mM Tris Base, 1mM EDTA Solution, 0.05% Tween 20, pH 9.0), Tris Base: 1.21 g, EDTA: 0.37 g, Distilled water: 1000 ml (100 ml for 10x stock). pH is set at 9.0 and then 0.5 ml of Tween 20 was added and mixed.

### Pre-processing, pre-staining and antibody staining of tissues for cycIF

Sample preparation for tissue imaging was performed using the following dewaxing, rehydration, pre-staining, staining protocol [13]. FFPE slides were incubated in a 60°C oven for 30 minutes. Paraffin was removed by immersing the slides in three changes of xylene for 5 minutes each. Slides were rehydrated in staining dishes containing 100% ethanol, 95% ethanol, 70% ethanol, 50% ethanol and then two changes of 1xPBS solution for 3 minutes each. Following rehydration, slides were placed in staining jars with Tri-Sodium citrate, pH 6.0, at the same time staining jars with distilled water and Tris-EDTA pH 9.0 buffer for antigen retrieval with HIER solutions were placed inside the steamer. Slides were steamed first with Tri-Sodium citrate, pH 6.0 for 20 minutes, then washed in hot distilled water and transferred to Tris-EDTA pH 9.0 buffer for 15 minutes. Slides were cooled to room temperature and washed three times with 1xPBS followed by blocking with Odyssey blocking buffer for 30 minutes using a moisture chamber (250-500uL per slide). Slides were then pre-stained by incubation with diluted secondary antibodies (1:2000) for 60 minutes. Slides were washed 3 times with 1xPBS, incubated with Hoechst 33342 (1ug/ml or 0.1uL/ml) in 250-500ul Odyssey blocking buffer for 15 minutes in a moisture chamber, and washed 3 times with 1xPBS in vertical staining jars. After imaging, slides were subjected to a round of fluorophore inactivation and washed 4 times in 1xPBS to remove residual inactivation solution.

### Staining with primary antibodies and Hoechst 33342

After slides were dewaxed and pre- stained, tissues were then stained in a moisture chamber by applying 250-500mL of diluted primary antibody (fluorophore-conjugated and unconjugated) on tissue followed by incubation at 4°C for 12 hours. All primary antibodies were diluted in Odyssey blocking buffer. Slides were washed 4 times in 1xPBS by dipping in a series of vertical staining jars. For Indirect immunofluorescence, slides were incubated in diluted secondary antibodies in a moisture chamber for an hour at room temperature. Slides were washed four times with 1xPBS, incubated in Hoechst 33342 at 1ug/ml in Odyssey blocking buffer for 10 minutes at room temperature and washed four times with 1xPBS. Slides were mounted prior image acquisition with 1xPBS containing 10% Glycerol (100ul per ml of 1xPBS), and coverslipped using a 24x60mm No.1 coverslip to prevent evaporation.

### Fluorophore inactivation (bleaching)

After imaging, fluorophores were inactivated by placing slides horizontally in 4.5% Hydrogen peroxide and 24 mM sodium hydroxide made in PBS for an hour at room temperature in the presence of white light. Following fluorophore inactivation, slides were washed four times with 1x PBS by dipping them in a series of vertical staining jars to remove residual inactivation solution. The detailed procedure is as follows: after image acquisition of each cycle, slides were placed in a vertical staining jar containing 1xPBS for at least 5 minutes. Coverslips were released from slides via gravity. The 100ml hydrogen peroxide stock solution was prepared with 15 ml of stock H2O2, 85ml of 1xPBS, 95mg of NaOH. Fluorophores were inactivated by placing slides horizontally in 4.5% H2O2 and 24mM of NaOH in PBS at room temperature in the presence of white light for an hour. Slides were imaged in successive immunofluorescence staining cycles.

### Image acquisition

Stained slides were imaged with a Zeiss Axio Scan Z1 slide scanning fluorescence microscope (ZEISS) using 20X objectives Plant-Apochromat 20x-0.8 M27). Colibri 7 camera (Zeiss Inc.) is used. Zen software (RareCyte Inc.) was used to sequentially scan the region of interest in four fluorescence channels. These channels are referred by the manufacturer as a: (i) DAPI channel with an excitation filter having a peak of 390 nm and half-width of 20 nm and an emission filter with a peak of 450 nm and half-width of 20 nm; (ii) EGFP channel having a 450-490 nm excitation filter and 500-550 nm emission filter (iii); Cy3 channel having a 538-562 nm excitation filter and 570-640 nm emission filter and (iv); Cy5 channel having a 625-655 nm excitation filter and 665-715nm; (v) Cy7 channel having a 720- 750nm excitation filter and 770-800 nm emission filter.

### Image processing and single cell protein quantification

Images with X20 magnification with pixel size equal to 0.33 µm are split using Zen software and uploaded onto OMERO [25]. Each CZI image has five channels and is converted to single channel-single plane tiff using bftools. An image was taken after each round of staining as well as each round of bleaching. Images of stained cycles have multiple Z planes while images of bleached cycles have one Z plane. Our comprehensive image analysis software (Crochet) for image processing, nuclear and cell segmentation, image quality control, downstream analysis was used for selection of most focused Z plane in stained images. Image registration, nuclear and cell segmentation, background subtraction and feature extraction were done using cycIFAAP software (https://pypi.org/project/cycIFAAP/) implemented in the Crochet image analysis suit with an embedded in-house trained Mask RCNN model for segmentation [26]. First, the nuclear boundaries are segmented based on DAPI signal. The cell boundaries are segmented using the distribution of E-Cadherin, PanKeratin, CD8, VEGFR and CD33 markers around nuclei. For cells not expressing any of the markers, the cellular boundaries are defined through expansion of nuclei boundaries until a neighbor cell is encountered or a maximum cell size is reached (3x nuclear size). Expression for each marker is extracted using their corresponding intrinsic sub-cellular expression zone. Crochet software is used for TMA core selection, exposure time and light intensity value normalization, correction of bleaching left-over signal, filtering non-specific binding sites, and removing cells that were lost through staining/bleaching cycles. Final corrected mean intensity values within single cell masks are taken from different zone localities within cell mask depending on biomarker intrinsic location (supp table with marker names and their intrinsic location). Expression values were median shifted to avoid taking log of small values and keep distributions close to the original marker distributions. Next, values are log2 normalized and Z scaled for all the following analysis. Manually curated threshold values are determined via careful inspection of images (supp table 4) for the purpose of defining marker positivity in cell type annotations. Threshold values are then used for annotating cell types using the hierarchical tree.

### Data analysis

All heatmaps are created using ComplexHeatmap package using Ward.B unsupervised hierarchical clustering method and Manhattan clustering distance calculation. All UMaps are created using Seurat package. A fixed minimum threshold value of zero is applied to log2 normalized and z scaled data. Linear dimensionality reduction is applied using all 39 protein markers and the UMaps neighbors is built with parameters data dimensionality 11, number of neighbors 30 and resolution 0.1 and further clustered using Louvain algorithm. Survival plots are created using R survminer and survival packages. All survival and t-test or Wilcoxon test p-values are FDR -Corrected using Benjamini-Hochberg procedure unless otherwise stated. Spatial interactions a are calculated using space software spatial score. Both marker-marker and cell-cell proximity is defined using a distance cut-off for neighboring cell/markers.

## Acknowledgement

We thank Gordon Mills for scientific discussions and critical reading of the manuscript, MDACC DQS-IT team for IT support, Guillaume Thibault for support with data analytics. This work is supported by grants from MDACC Support Grant P30 CA016672, funding from the MDACC Colorectal Cancer Moon Shot program, U01 CA253472. This work was supported by the Kavanagh Family Foundation and Kevin T. Doner Memorial Fund (no grant number, philanthropic).

## Author contributions

Conceptualization and study design: AK, MO. Data generation: ZD, MS. Quantitative analysis: BB, AK. Analysis and interpretation: AK, BB, ZD, MO, JW, HW, GMB, NH. Sample curation: HW, MO, ZD. Writing: AK, MO, ZD, BB, JW, GMB, NH. Funding Acquisition: AK, GMB, MO. Resources: AK, MO. Project administration: AK, MA. Data quality control: AK, MO, HW.

## Data sharing statement

All raw images are available in the OMERO database of Korkut laboratory at MDACC. The spatial feature maps are shared on the Korkut laboratory OMERO resource. The integrated analysis script to reproduce the results in this manuscript is available. The processed spatial expression within supplementary data. The meta data for antibody information, assays and imaging parameters are shared in supplementary tables.

